# Emergence of potato virus Y outbreaks in tomatoes in Brazil, the disease and spread

**DOI:** 10.1101/2024.05.17.594728

**Authors:** Vivian S. Lucena, Erich Y. T. Nakasu, José L. Pereira, Camila M. Rêgo-Machado, Cristiano S. Rodrigues, Bernardo Ueno, Ivair J. Morais, Alice K. Inoue-Nagata

## Abstract

The emergence of ’Mexican Fire’ disease in Brazilian tomato fields, attributed to potato virus Y (PVY), has raised concerns. Characterized by severe necrosis on median leaves, the definitive etiological agent of this disease remained unverified despite PVY detection in symptomatic plants. Our study aimed to elucidate the causal agent, occurrence, spread, and symptomatology of Mexican Fire. Deep sequencing of tomato leaves with typical necrotic symptoms confirmed the association with PVY, reinforcing its role as the causal agent. Serological tests with a PVY-specific polyclonal antibody consistently correlated symptoms with virus presence in a fresh market tomato field, with higher PVY incidence near older tomato and maize plants. Necrotic leaf distribution analysis revealed a predominant occurrence in median leaves, progressing upwards. Deep sequencing of symptomatic field samples exclusively detected PVY, reaffirming its role in symptom induction. Importantly, PVY inoculation under field and greenhouse conditions fulfilled Koch’s Postulates, triggering leaf necrosis. Our findings unequivocally establish PVY as the causal agent of Mexican Fire disease, shedding light on its etiology, incidence, spread, and symptom expression, crucial for effective disease management strategies.

## Introduction

Potato virus Y (PVY) is classified in the species *Potato virus Y*, a member of the genus *Potyvirus* in the family *Potyviridae*. The PVY genome consists of a monopartite single-stranded RNA enclosed by coat protein subunits, forming a flexuous filamentous particle (Inoue-Nagata et al. 2022). Transmission occurs naturally by a diverse array of aphid vector species in a non-persistent manner (Davie et al. 2017; Rizk et al. 2020). PVY exhibits a wide range of plant hosts, including agricultural relevant species such as tomato, potato, peppers, and tobacco, as well as numerous weed species (Edwardson and Christie 1997; Jeffries 1998).

Historically, PVY has been most important in potato plants, mainly because of the limited availability of resistant varieties. However, there is a growing concern that PVY is emerging as a significant threat to tomato plants as well (Hasiów-Jaroszewska et al. 2015; Sihelská et al. 2016; Grbin et al. 2023). Worldwide, PVY-infected tomato plants present a wide range of symptoms, including stunting, necrotic leaf lesions, and distinctive features such as leaf wrinkling and mosaic patterns (Rosner et al. 2000; Aramburu et al. 2006; Hasiów-Jaroszewska et al. 2015; Chikh-Ali et al. 2016).

In the past, PVY posed a major threat to tomato production in Brazil, but its prevalence declined sharply after the introduction of resistant cultivars by the late 1960s (Nagai and Costa 1969; Nagai et al. 1992). As a result, currently tomato growers seem to have limited concern regarding the presence of PVY-resistant traits.

In recent years, a new disease named ‘Mexican Fire’ (Fogo Mexicano) has emerged, affecting tomato plants in various regions of Brazil, including the South, Southeast, and Central-West areas. Growers have reported the sudden appearance of foliar necrotic lesions, which can quickly lead to the complete loss of photosynthetic areas in the plants. Mostly, the necrosis is observed in the median leaves, and less frequently in older and younger leaves. Remarkably, these symptoms consistently coincide with the detection of PVY through immunological and molecular assays (unpublished observations). Although necrosis has occasionally been associated with PVY infection in other countries (Gebre-Selassie et al. 1987; Rosner et al. 2000; Aramburu et al. 2006; Mascia et al. 2010), confirming PVY as the definitive cause of Mexican Fire in Brazil remains challenging due to unsuccessful attempts to reproduce these symptoms via inoculation tests (unpublished results). This has thus far prevented the fulfillment of Koch’s Postulates.

Given the increasing reports of severe necrotic symptoms associated with PVY in key tomato-growing areas of Brazil and the lack of thorough information on the causes, symptoms, prevalence, and spread of the disease, this study aims to provide a detailed insight into Mexican Fire disease. Using a range of methods, we investigated whether PVY is solely responsible for causing the disease’s symptoms, both in natural conditions and controlled experiments. We rigorously examined the relationship between PVY infection and symptom development, aiming to confirm PVY as the primary cause of Mexican Fire disease and contribute to a better understanding of how to manage its impact on tomato crops.

## Material and methods

### High throughput sequencing (HTS) of necrotic plants

We analyzed the viral populations in three composite leaf samples collected from tomato plants with necrotic leaf symptoms in the middle of the plant and without apparent symptoms in the upper or lower leaves. Fungal or bacterial etiology was ruled out based on diagnostic testing of the presence of bacterial flow and microscopic observation of fungal structures following incubation in humid chambers. The first sample, referred to as RNY1, consisted of a pool of leaves from plants collected in several Salad type tomato fields in Pelotas, Rio Grande do Sul state, in 2014. The second sample, RNY2, was composed by a pool of tomato cv. Dominador samples collected in Araguari, Minas Gerais state, in 2014. Partial results generated to RNY1 and RNY2 were published by Martins et al. (2021). The third sample, referred to as TOMNEC, consisted of a pool of leaves from a single tomato cv. Guará plant collected in Taquara, Federal District, in 2019, in the monitored field described below. This plant was tested negative by ELISA using polyclonal antibodies for other tomato-infecting viruses: groundnut ringspot virus (GRSV), pepper yellow mosaic virus (PepYMV), and tomato mosaic virus (ToMV).

In order to determine the viruses present in the tomato samples, total RNA was extracted using RNeasy (Qiagen) following the manufacturer’s instructions. Library preparation was carried out after the removal of ribosomal RNA from plants with TruSeq Stranded Total RNA LT Sample Prep Kit. Sequencing of 100 nt PE was performed on an Illumina Novaseq Platform at Macrogen, Inc. (Seoul, South Korea). Reads were trimmed with Trimmomatic 0.35, set to an automatic Phred score to 0.34. The contigs were assembled using the Velvet algorithm v. 1.2.10 (Zerbino and Birney 2008). Contigs were analyzed using Megablast and tBLASTx on Geneious 9.1.8 with a virus RefSeq library (downloaded on 02/18/2019). The contigs of interest were used as a reference for the assembly of the complete PVY genome using the Geneious mapper (up to 25 iterations). The complete genome sequence was analyzed using the BLASTN tool for comparison with other database sequences (https://blast.ncbi.nlm.nih.gov/Blast.cgi).

### Monitoring plants in a field for detecting natural infection by PVY

Plants were monitored for viral infection in a small farm located in Taquara in the greenbelt of Brasília (coordinates: -15.659358, -47.534959, Figure 1), Federal District, Brazil. Three areas, referred as 1, 2 and 3, within the farm were weekly monitored soon after transplanting for a period of 12 weeks. Plants of the Area 1 and Area 2 were transplanted at February 8, 2019 (cultivars BS12, Santy and Guará), and Area 3 at March 18, 2019 (cultivars Santy and Guará). Other crops were present in the surrounding area, such as tomato, sweet pepper, zucchini, cucumber, cauliflower and cabbage (Fig. 1).

**Fig 1.**
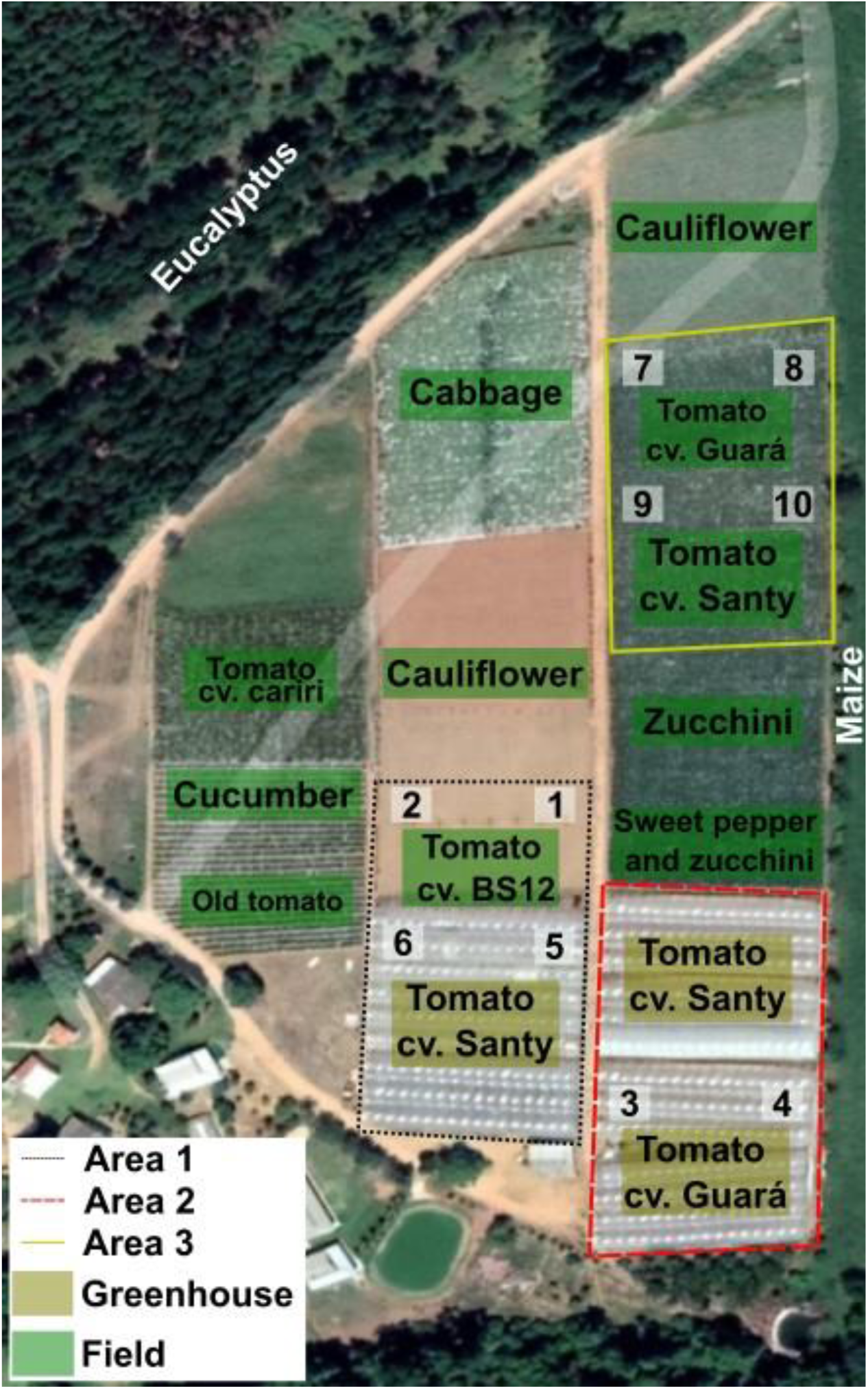
Schematic representation of the monitored tomato field derived from a Google Maps satellite image (accessible at https://www.google.com/maps). The areas of interest for sampling are delineated in Areas 1, 2, and 3, identified by the black dotted, red dashed, and yellow solid lines, respectively. The green background denotes open-field cultivation, while the golden color indicates areas under protected cultivation. Collection sites for plots 1 to 10 are represented by numbered squares, facilitating the identification of study areas.

Ten plots of 25 plants each (five planting rows and five plants per row) were delimited close to the borders of each area (plots 1 to 10, Fig. 1). Three cultivars were monitored: BS12 (BlueSeeds) (plots 1, 2), Guará (H.M. Clause) (plots 3, 4, 7, 8), and Santy (Sakata Seeds Sudamerica) (plots 5, 6, 9, 10). Plot 6 was closer to an old tomato field in the final stage of harvest. These tomato plants were removed ∼6 weeks after transplanting in Area 1. The presence of PVY-infected plants in this old tomato field was confirmed by serology in plants with strong necrosis in the median leaves (not shown). Plots 4, 8, and 10 were closer to a maize field planted beside the evaluated areas. Plots 3, 4, 5, and 6 were covered with a plastic roof, but open laterally.

The plants were examined once a week, the symptoms registered, and young leaves collected from each monitored plant. Necrotic spots in the abaxial and adaxial median to top leaves, and mild mottling on upper leaves were considered as the typical Mexican Fire symptoms. Plants showing typical symptoms of tospovirus infection, such as strong necrosis only on upper leaves, were clearly discerned from those with necrosis in the median portion of the plant. A virus detection test was employed in these collected samples, consisting of antigen-trapped indirect ELISA using antibodies against the following viruses: GRSV, ToMV, PepYMV, and PVY. These are the typical RNA viruses found in tomato plants in Brazil (Inoue-Nagata et al. 2016).

### Correlation between PVY detection and necrotic leaf symptoms

In order to evaluate the correlation of the occurrence of necrotic symptoms with PVY infection, 249 samples were randomly collected, from 138 plants showing necrosis in the abaxial (in some cases also in the adaxial) part of the leaf, which are typical of Mexican Fire disease, and from 111 asymptomatic plants. All samples were tested for the presence of PVY by ELISA.

### Distribution of necrotic symptoms and detection of PVY in the leaves of symptomatic plants

In order to investigate the distribution of the necrotic symptoms throughout the adult plants, visual analyses were carried out on 10 plants of cultivar Guará and 11 of cultivar Santy previously confirmed by serology to be infected with PVY. Each leaf was evaluated from the top to the bottom (oldest leaves) for the presence of leaf necrosis. Then, three plants of cultivar Santy were collected, all the leaves removed and tested by ELISA to determine which leaves contained virus particles detectable by serology.

### Evaluation of the spread of PVY in the field

The incidence rate and distribution pattern of PVY symptomatic plants were analyzed in the Areas 1, 2 and 3 (Fig. 1). All plants (inside and outside the plots) were visually evaluated for the presence of necrotic symptoms, typical of Mexican Fire disease. This analysis was performed three times (6, 8 and 10 weeks after transplanting - WAT) in Area 1 and Area 2, and five times (9, 11, 13, 15 and 17 WAT) in Area 3. The incidence rate was calculated for each time point in the three areas. Then, at 10 WAT for Areas 1 and 2, and at 13 WAT for area 3, the incidence rate was determined for each cultivar according to the distance from the border. Excluding four rows of both borders, the planting row of 50m was divided in 10 blocks (1 to 10) of 5m, and the symptomatic plants were counted in the block.

### Inoculation of isolates of PVY in field and greenhouse cultivated plants to confirm the production of necrotic symptoms in infected plants

The PVY-TOMNEC isolate was used to demonstrate whether PVY causes the typical expression of necrotic symptoms in median leaves of tomato plants. Two experiments were conducted, the first in the field and the second in a greenhouse. For the field experiment, PVY-TOMNEC was used to inoculate 12 plants of each of the cultivars Santy and Guará. Plants were 10 weeks old, and were sap inoculated twice (on May 20 and May 27, 2019) with an extract prepared with 0.05M sodium phosphate buffer, pH 7.5. The same number of plants were mock inoculated with inoculation buffer. Before inoculation, the plants were tested by ELISA to confirm they were not infected by PVY. Plants were kept in the field for symptom development and after 15-22 days, samples were collected for ELISA against PVY.

The second experiment was conducted in the greenhouse. Five to nine plants each of cultivar Santy and Guará were inoculated at 15 and 30 days after transplanting with two isolates: PVY-TOMNEC and PVY-363. The isolate PVY-363 was identified in tomato with necrotic symptoms and is part of our virus collection. Three inoculations were performed in the interval of 7 days. ELISA was performed for PVY detection 30 days after the first inoculation. Plants were weekly examined for symptom development.

### Phylogeny

For phylogenetic analysis of the PVY sequences, we expanded our dataset by including 25 representative sequences obtained from various strains and countries. To ensure accurate alignment of these sequences, the Mafft algorithm (Katoh 2002) was employed. The iqtree2 software (Minh et al. 2020) facilitated the construction of the phylogeny relationship among these samples using ML-method. ModelFinder (Kalyaanamoorthy et al. 2017) aided in selecting the optimal model of sequence evolution. To enhance the reliability of our analysis, we subjected it to 10,000 bootstrap replicates. Finally, the tree was edited using iTOL (Letunic and Bork 2021).

## Results

### HTS for identification of the causal agent of Mexican Fire

Two pools of leaf samples collected from plants presenting Mexican Fire symptoms were first analyzed by HTS to investigate which viruses were associated with the necrosis: RNY1 from Rio Grande do Sul, the southernmost state of Brazil; and RNY2 from one of the largest tomato growers for fresh market in Brazil, located in Araguari, Minas Gerais state. The following virus sequences were identified in the RNY1 library: the potyviruses PVY and potato virus P (PVP), and the polerovirus potato leafroll virus (PLRV). RNY2 contained sequences of the potyviruses PVY and pepper yellow mosaic virus (PepYMV), the crinivirus tomato chlorosis virus (ToCV), the amalgavirus Southern tomato virus (STV), the tospoviruses tomato chlorotic spot virus (TCSV) and groundnut ringspot virus (GRSV), the tobravirus pepper ringspot virus (PepRSV), the tobamovirus pepper mild mottle virus (PMMoV), the tymovirus tomato blistering mosaic virus (ToBMV), the potexvirus potato virus X (PVX), and the cucumovirus cucumber mosaic virus (CMV) (Martins et al. 2021). PVY was the only virus detected in both libraries, suggesting that PVY is the sole causal agent of the necrosis symptom in the median leaves of those plants.

To make sure that the necrosis symptoms were caused solely by PVY, one plant with typical necrotic symptom of Mexican Fire was also selected for deep sequencing. The isolate TOMNEC from cv. Guará (Fig 2H) was collected in the monitored field (Area 3), and the RNA was sequenced. 6,800,000 viral reads were detected in the library, and all corresponded to the PVY genome, confirming that this plant was infected solely by PVY.

**Fig 2.**
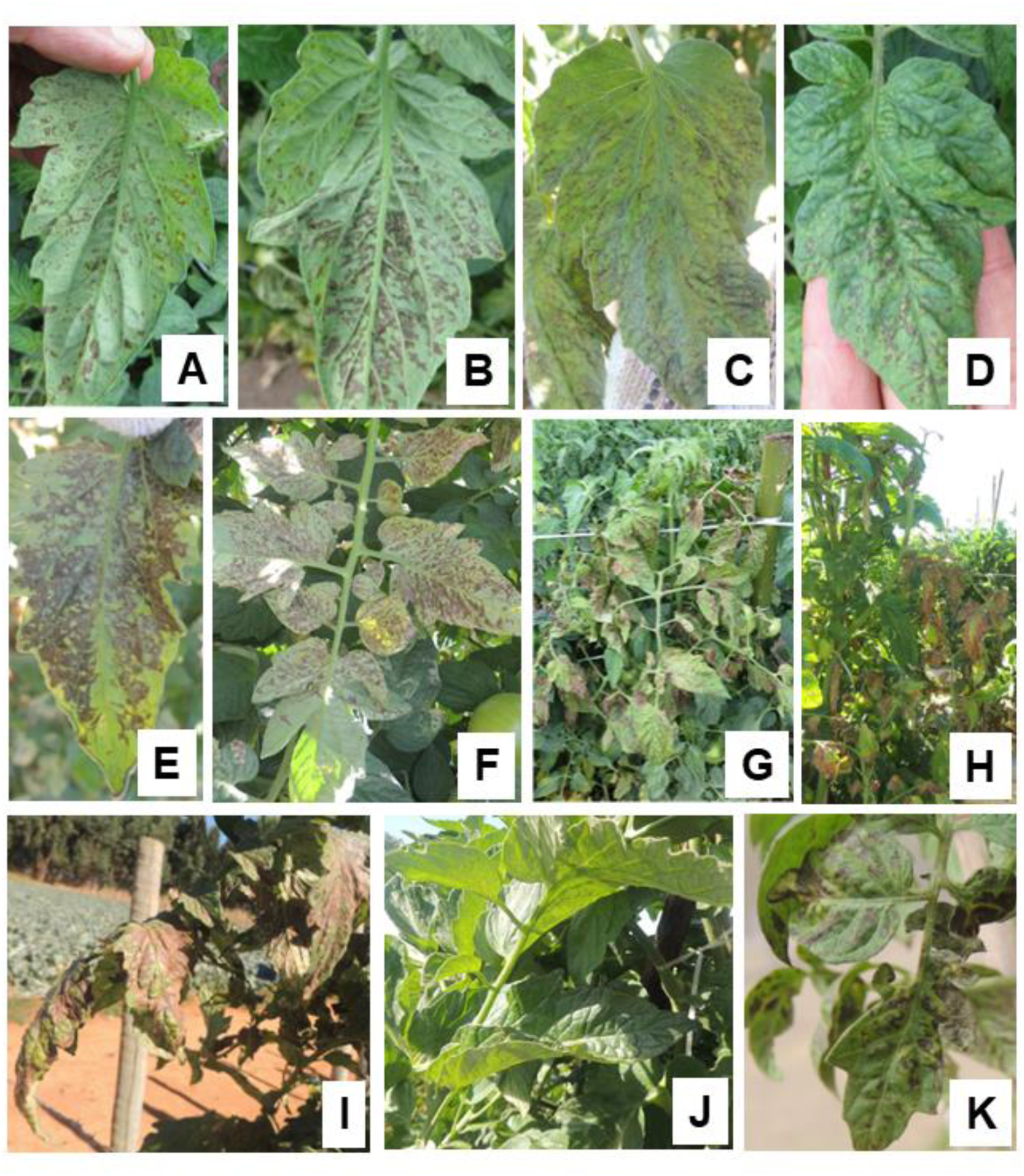
Tomato plants infected with potato virus Y (PVY) exhibit variations in necrotic symptoms. Initially, necrotic spots are observed on the abaxial side of the leaves (A, B), followed by their appearance on the adaxial side (C, D). Subsequently, these necrotic lesions progress, eventually covering substantial portions of the plant (E-G). The PVY TOMNEC isolate was obtained from a plant in the field displaying necrotic spots (H) and was confirmed to be exclusively infected with PVY. Koch’s Postulates were demonstrated both in the field (I, inoculated and infected; J, mock-inoculated and non-infected) and in the greenhouse (K).

The sequence of PVY-TOMNEC (GenBank accession OR497354, 9,689 nucleotides), PVY-RNY1 (OR497355, 9,687 nucleotides) and PVY-RNY2 (OR497356, 9,688 nucleotides) had a few nucleotides missing in the 5’ terminal end, and the RNY1 sequence in the 3’ end. The sequences share 91.3-91.5% nucleotide identity with the C strain of PVY isolate PRI-509 (EU563512, 100% coverage and Evalue of 0.0). They share within each other 95.2-98.2% nucleotide identity, being PVY-RNY1 and PVY-RNY2 the closest ones with 98.2% nucleotide identity.

A phylogenetic analysis was performed after the alignment of PVY-RNY1, PVY-RNY2, PVY-TOMNEC and 25 PVY complete genome sequences retrieved from public databases, most of them from pepper and tomato plants. The three tomato PVY isolates from Brazil were tightly clustered, indicating a close relationship and a common origin (Fig 3). They were grouped in a high bootstrap support with all sequences derived from pepper plants (4 isolates), as well as with all the six isolates classified as C strain, from tomato and potato plants. This strongly supports that the Brazilian tomato isolates share a common ancestor with most isolates of the C strain that are able to infect pepper, tomato and potato plants. The other isolates were highly variable and consisted of tomato and potato isolates, from O, N and other variants.

**Fig 3.**
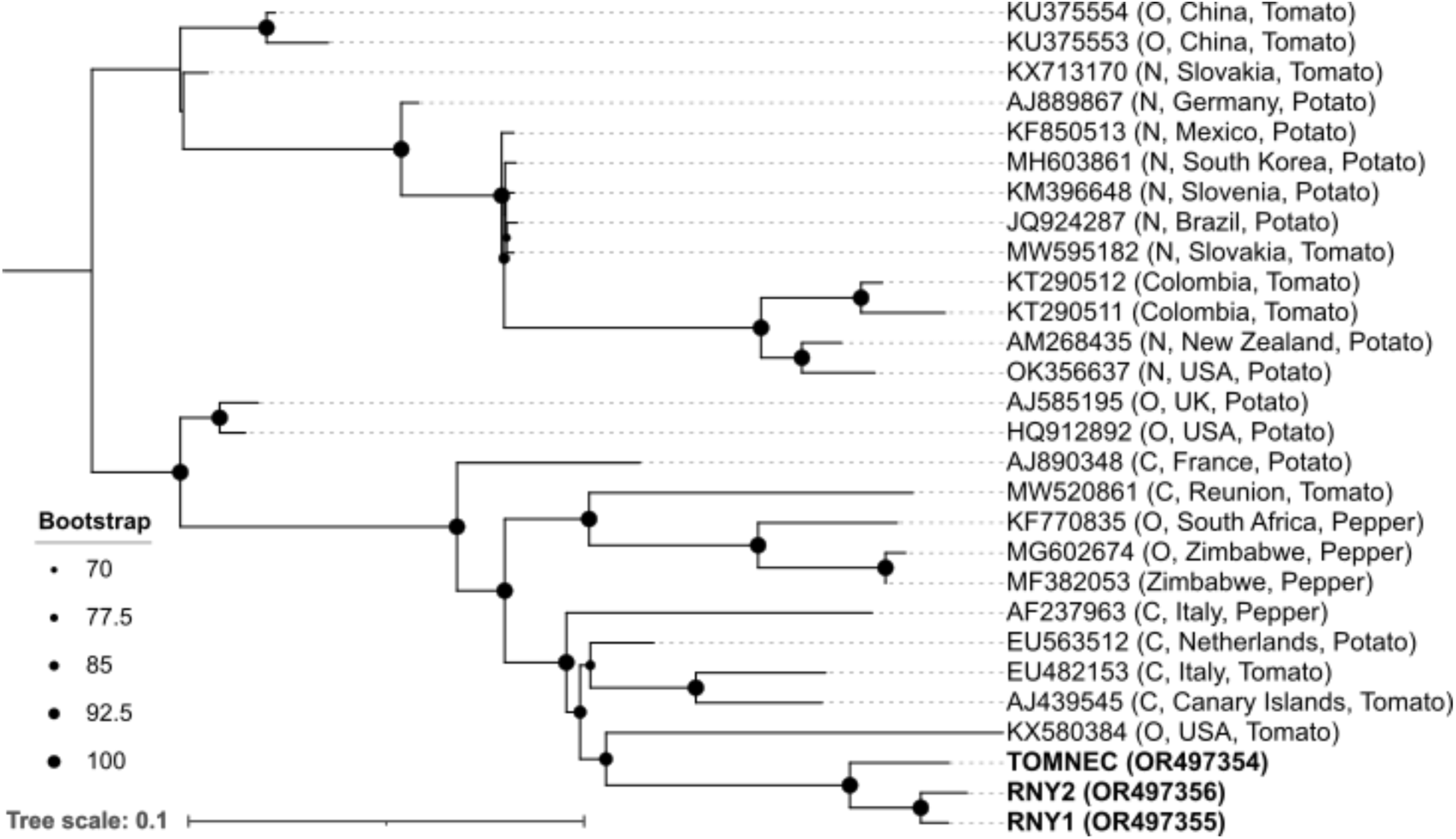
ML-Phylogenetic tree of three Brazilian tomato PVY near complete genome sequences with 25 representative PVY sequences. Alignment was performed using Mafft algorithm. The sequences are identified with the GenBank accession number, followed by the strain classification, country, and host, when available, between brackets. The Brazilian isolates are highlighted in bold letters, with the corresponding accession numbers. The tree was edited using iTOL webserver tool. Bootstrap values are represented by the size of black circles at the nodes of branches. PVY sequences determined in this study are indicated by bold letters. Bar indicates substitutions per site.

### Association between necrotic symptom and PVY infection

Next, we investigated if PVY was consistently associated with plants with necrosis in the median leaves. From a commercial fresh market tomato field, plants were inspected for the presence of necrosis symptoms, and samples were collected and tested by serology for PVY detection. A total of 249 leaf samples were collected from plants with and without necrosis symptoms at 67 days after transplanting. PVY was detected in 129 out of 138 symptomatic plants (93.4%). The nine negative plants were possibly infected with other pathogens that cause necrosis, such as GRSV and *Stemphylium* sp., both present in the area. From the 111 asymptomatic plants, 16 (14.4%) were positive for PVY detection, and the remaining 95 (85.6%) were negative. The strong association between viral presence and symptoms in the plant population was evident, with an odds ratio of approximately 104.27, indicating that the virus is significantly more likely to be found in symptomatic plants compared to asymptomatic ones.

### Monitoring tomato plants for PVY infection and its association with necrosis symptoms: symptom development, distribution and spread

Three tomato fields were weekly monitored for the detection of PVY and the appearance of plants with necrosis symptoms. We aimed to understand whether all PVY-infected plants developed necrotic symptoms, and if all plants with necrosis were infected with PVY. Ten plots with 25 plants each were delimited in an area where the necrotic disease often occurs. The first PVY infected plant, detected by serology, was observed 3 WAT in plot 4, which is close to the maize field (Fig 1). In this plant, necrotic lesions appeared in the abaxial part 5 WAT, i.e., 2 weeks after PVY detection by serology. During the monitored period of 12 weeks, PVY was detected in 30 symptomatic plants (12%, 30/250 – Table 1). Furthermore, the virus could be detected in 13 asymptomatic plants, all of them in the last evaluation (Table 1). Plot 6 presented the greatest number of infected and symptomatic plants, in a total of 7 symptomatic (28%) and 10 ELISA positive plants (40%) (Table 1). This plot was located beside an old tomato field, which was eliminated 6 WAT of plants in this monitored field (Fig 1, Table 1). Plots closer to maize (plots 4, 8, 10) and old tomato (plot 6) presented 70% of the symptomatic plants and 67.4% of infected plants, suggesting that these crops might act as reservoirs of virus and/or aphid vectors (Fig 1).

**Table 1.** Plants with necrotic symptoms in the monitored plots after 12 weeks.

| Plot<sup>1</sup> | Cultivar | Closest crops to the plot<sup>2</sup> | No. symptomatic plants<sup>3</sup> | No. PVY positive plants by serology<sup>3</sup> |
| --- | --- | --- | --- | --- |
| 1 | BS12 | Zucchini | 1 (4%) | 2 (8%) |
| 2 | BS12 | Cucumber | 0 (0%) | 0 (0%) |
| 3 | Guará | Tomato cv. Santy | 1 (4%) | 2 (8%) |
| 4 | Guará | Maize | 4 (16%) | 6 (24%) |
| 5 | Santy | Tomato cv. Santy | 1 (4%) | 1 (4%) |
| 6 | Santy | Old tomato | 7 (28%) | 10 (40%) |
| 7 | Guará | Cabbage | 2 (8%) | 3 (12%) |
| 8 | Guará | Maize | 5 (20%) | 7 (28%) |
| 9 | Santy | Cabbage/cauliflower | 4 (16%) | 6 (24%) |
| 10 | Santy | Maize | 5 (20%) | 6 (24%) |
| **Total** |  |  | **30 (12%)** | **43 (17%)** |

<sup>1</sup>Plots were located at the border of the field.
<sup>2</sup>Other crops were cultivated close to the evaluated plots.
<sup>3</sup>Percentage between brackets.

The plants tested negative for the tomato-infecting viruses PepYMV, ToMV, and GRSV, as confirmed by ELISA assays. However, tospovirus infection symptoms were observed in some plants in the field outside the plots (data not shown).

As all plants were weekly evaluated for the presence of necrotic symptoms, and tested for PVY infection, we investigated the time span for symptom expression after infection. Most of the plants showed visible symptoms 2 weeks after infection (i.e., positive detection by serology), but some took up to 5 weeks. For 13 plants, the necrotic symptoms were not observed, most likely because the virus was detected in the last weeks of evaluation and the symptoms would develop later. This again confirmed that the presence of PVY was strongly associated with the appearance of the necrotic symptoms.

Next, in order to confirm if the necrosis occurs preferentially in median leaves, mature and infected plants (with 23-26 leaves) were randomly selected from each cultivar, Santy (10 plants) and Guará (11 plants), to evaluate the position of the necrotic leaves in the entire plant. The necrotic symptoms were observed mostly from the 3rd to the 14th leaf, confirming that the symptoms were present in the median part of the plant. Then, we investigated if the virus could be detected in asymptomatic leaves in infected plants. Three plants of the cv. Santy were selected for symptom evaluation and virus detection by ELISA. The evaluated plants had 17-21 leaves, and the necrotic symptoms were observed in leaves 3-13, while the virus was detected in leaves 1-16 (Supplementary Table 1). The chi-square tests revealed varying associations between symptoms and ELISA results for the three plants. Plant 1 exhibited no statistically significant association, with a chi-square statistic of 1.987 and a *p*-value of approximately 0.107 (*p* > 0.05). In contrast, both Plant 2 and Plant 3 displayed significant associations, with chi-square statistics of 2.59 and associated *p*-values of 0.003 and 0.01, respectively (p < 0.05). These findings suggest that symptoms and ELISA results are significantly associated in Plant 2 and Plant 3, while no such association was observed in Plant 1. Taking together, the results indicates that, except for the old leaves, the virus was widespread within the plant but the symptoms were prevalent in the mid portion.

The symptoms started as small necrotic lesions in the abaxial part of the leaflets in the median portion of the plant (Fig 2A, 2B). At this stage, the symptoms were not visible in the adaxial part. Later, the lesions coalesced and formed larger lesions in the interveinal part, visible in the adaxial part, and covering the entire leaf (Fig 2C-2G). Then, the necrotic symptoms were gradually being observed in younger leaves, including those of the top. During cold days, infected plants were more chlorotic than healthy plants (not shown).

### Distribution and spread of diseased plants in the field

For assessing incidence, and understanding the spatial distribution of diseased plants, we focused on the surveillance of symptomatic plants within the three fields: Area 1 to Area 3 (Fig.1). Area 1 and Area 2 encompassed plants of cultivars Santy and Guará, covered by plastic for rain protection, alongside a smaller open field area cultivated with the BS12 cultivar, comprising a total of approximately 5,700 plants.

Over the course of our study, three evaluations, spaced two weeks apart, were performed, resulting in the identification and spatial distribution of symptomatic plants.

In the case of Area 1, the initial observation of tomato plants displaying characteristic necrotic symptoms occurred 6 WAT. Within this region, 0.4% of the plants exhibited symptoms, and their distribution appeared random. Subsequently, within a two-week interval, the incidence of symptomatic plants increased to 1.3%, with a notable concentration along the borders, particularly in proximity to the pre-existing tomato plants. Then, after an additional two weeks, symptomatic plants reached an incidence rate of 5.7%, again with a pronounced concentration along the borders.

In Area 2, the initial evaluation at 6 WAT revealed that only 0.3% of the plants exhibited necrotic symptoms, with a higher occurrence near the field borders, adjacent to the maize plants. In subsequent evaluations, the incidence rate displayed slight increments, reaching 0.5% in the second assessment and 1.7% in the third evaluation. Likewise, the preponderance of symptomatic plants was concentrated near the field border, in proximity to the maize field. Notably, the symptomatic plants exhibited clustering tendencies, although isolated instances were also observed. Within this area, infection onset initiated at a later stage, and damage was likely not pronounced.

In Area 3, the plant population consisted of cv. Guará and Santy, which were situated within an open field (Fig. 1). These plants underwent regular evaluation at two-week intervals, in a total of five evaluations. The study encompassed a total of ∼14,450 plants, organized into 84 lanes. During the initial assessment (9 WAT), only 0.1% of the plants exhibited necrotic symptoms. However, in the subsequent evaluation, the incidence of symptomatic plants increased to 1.7%, with a pronounced concentration along the field’s border, in close proximity to the maize plants. Subsequent evaluations showed a steady escalation in incidence rates, reaching 5.4%, 13.2%, and in the final evaluation at 19.0%. As the evaluations were done visually, we expect that more plants were effectively infected.

It is important to note that plants displaying symptoms were distributed across the entire field, but their density was likewise higher along the field border and closer to the maize field. In view of the potential border effect, we quantified the distribution of PVY-infected plants along the planting rows in our study area. To mitigate interference, the analysis excluded the four rows at the border. Each row was then divided into 10 blocks, each spanning 5m and all rows except four at each border. Symptomatic plants were counted within each block, and the percentage was plotted on a graph, with the predominant crop adjacent to the field highlighted (Fig. 4). This analysis was conducted for each cultivar in the study area, resulting in five datasets: two for plants of cv. Guará and three for cv. Santy. The graphical representation revealed a notable trend: the incidence of symptomatic plants was highest near the border adjacent to an old tomato field (Fig. 4A), as well as near maize fields (Fig. 4B-E). This pattern was in sharp contrast to the interior portion of the field and the borders adjacent to other crops. These findings strongly suggest that the old tomato field acted as a source of PVY for the new tomato field. Additionally, the maize field appeared to serve as a significant reservoir or source of the vector, contributing to the spread and propagation of the disease among the plants.

**Fig 4.**
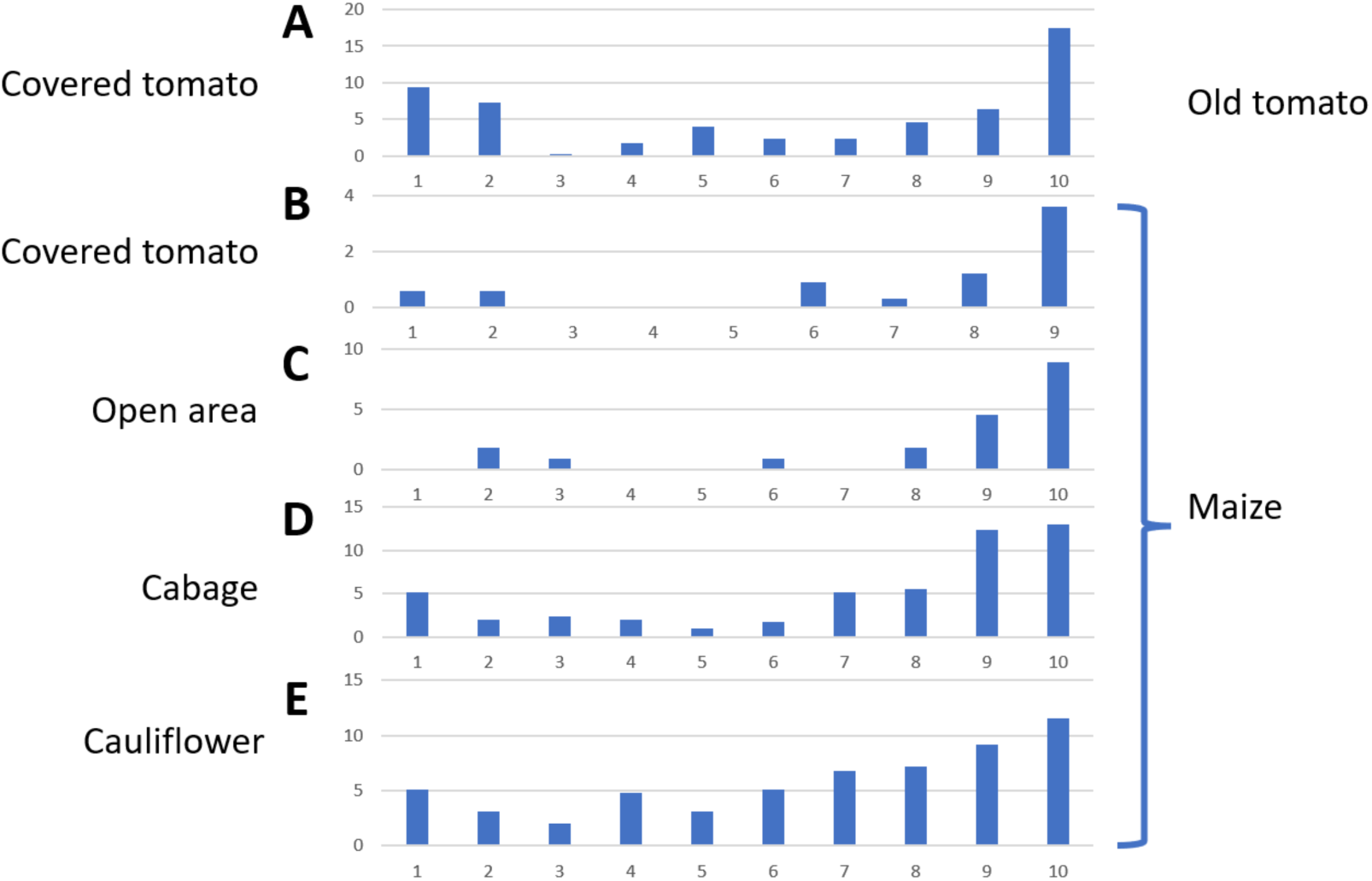
Graphical representation of the incidence of symptomatic plants along the planting rows. Each bar represents a 5-meter segment, with a total of 10 bars corresponding to one row with 50 meters and ∼86 plants. The number of symptomatic plants was counted within each segment, with percentages plotted accordingly. The predominant crop adjacent to the field is highlighted for reference. Cultivars are indicated as follows: A. Cultivar Santy, 26 rows; B. Cultivar Santy, 39 rows; C. Cultivar Guará, 13 rows; D. Cultivar Guará, 34 rows; E. Cultivar Santy, 34 rows.

### Inoculation of PVY in tomato plants

All results indicate that PVY is the causal agent of the Mexican Fire disease. To confirm its etiology, two inoculation experiments were performed. The first trial was carried out to test if plants cultivated in greenhouse conditions respond to infection with the necrotic symptom, when inoculated at 15 and 30 days after transplanting. Two PVY isolates were used, PVY-TOMNEC and PVY-363, both from tomato, in two cultivars, Santy and Guará. The seedlings were transplanted in pots and kept in a greenhouse throughout the trial. Five to nine plants were used for each test. Necrotic lesions were observed in both cultivars (Fig. 2K) and in both inoculation times, whereas the rate was lower in plants inoculated at 30 days after transplanting (Table 2). Plants mock-inoculated were not infected, and no necrotic symptoms were observed. This result confirmed that the infection of PVY caused necrotic spots in tomato cv. Santy and Guará.

**Table 2.** Plants with necrotic symptoms after inoculation with two isolates of PVY.

| Isolate | Inoculation time<sup>1</sup> | Cultivar |  |
| --- | --- | --- | --- |
|  |  | Santy<sup>2</sup> | Guará<sup>2</sup> |
| PVY-TOMNEC | 15 DAT | 4/5 | 3/5 |
|  | 30 DAT | 2/7 | 3/9 |
| PVY-363 | 15 DAT | 3/5 | 3/5 |
|  | 30 DAT | 0/8 | 1/7 |
| Mock inoculated | 15 DAT | 0/5 | 0/5 |
|  | 30 DAT | 0/5 | 0/5 |

<sup>1</sup>Plants were inoculated 15 or 30 days after inoculation (DAT).
<sup>2</sup>Number of infected plants with necrotic symptoms/number of inoculated plants at 30 days after inoculation.

Then, plants of the monitored field were inoculated. Twenty-four asymptomatic and uninfected plants (as tested by ELISA) of each cultivar, Guará and Santy, were selected 9 WAT. Twelve plants from each cultivar were mock inoculated with inoculation buffer, and 12 with a fresh leaf extract from PVY-TOMNEC. Two weeks after inoculation, all 12 plants of cv. Santy became infected as confirmed by serology and 10 of those showed the typical necrotic symptoms in the leaves (Figure 2I). For cv. Guará, 10 out of 12 were positive for PVY infection, and all of them showed the typical necrotic symptoms 4 weeks after inoculation. Taken together, these results confirm that PVY cause the necrotic symptoms characteristic of the Mexican Fire disease.

## Discussion

The earliest documented occurrence of PVY on tomatoes in Brazil dates to 1956 when Silberschmidt and Tommasi (1956) detailed the observation of symptomatic plants collected in the São Paulo state (Piedade county). These plants exhibited reduced leaf size, rugosity, narrow and thin leaves, as well as vein clearing and thickening. Subsequently, Costa et al. (1960) reported tomatoes displaying ’Risca do Tomateiro’ (Tomato Streak) symptoms due to PVY infection, characterized by smaller leaf size, epinasty, streaks on mid-leaves, and mild mosaic manifestations in young leaves. The term ’Risca’ denotes streaks occurring within or parallel to the leaf veins, primarily on the abaxial side of the leaves (Costa et al., 1960).

The introduction of the first resistant variety, named Ângela, occurred in 1969 through the crossbreeding of the resistant PI 126410 (*Solanum lycopersicum* x *S. pimpinellifolium*) with the susceptible commercial variety Santa Cruz (Nagai and Costa 1969). This variety, along with its derived selection lines, was widely used in roughly 70% of tomato fields for fresh market (Nagai, 1993), thereby contributing significantly to the effective management of Risca do Tomateiro. Presently, the newly cultivated tomato varieties in Brazil are exclusively hybrids, with no available information regarding their resistance to PVY.

In our monitored experimental plots, the initial detection of PVY-infected plants occurred approximately 30 days after transplanting, with the corresponding symptom manifestation occurring two weeks afterwards. Throughout this interim, aphids were not visibly present on the plants during routine inspections; however, winged aphids were captured using yellow traps (data not presented). This observation confirms the presence of vectors within the vicinity.

Plots in proximity to the former tomato cultivation area (plot 6: 7 infected plants per plot) and maize plants (plots 4, 8, 10: average of 4.7 infected plants per plot) exhibited a higher incidence of infection compared to those closer to other crops (plots 1, 2, 3, 7, 9: average of 1.6 infected plants per plot). It is noteworthy that plot 5, situated close to tomato plants of similar age, harbored only one infected plant. This observation strongly supports the hypothesis that established tomato plants are reservoirs of PVY for newly planted tomatoes, while maize serves as a source of aphids. This result highlights the significance of planting tomatoes at a considerable distance from host crops of PVY and aphids.

The symptoms induced by PVY on tomatoes are highly variable, according to the specific isolate and the tomato variety under consideration (Aramburu et al. 2006; Zitter, 2014). These symptoms include the common mosaic and mottling (Morel et al. 2000), along with symptomless infection, mosaic, necrosis, yellowing, leaf deformation and stunted growth (Aramburu et al. 2006). Usually, PVY elicits faint mottling and mild distortion, and also more severe manifestations such as necrosis and leaf distortion, flower necrosis, and irregular fruit ripening (Zitter, 2014). Rosner et al. (2000) described PVY isolates able to induce necrotic symptoms such as those of Mexican Fire in Israel, suggesting that these isolates are unique, and distinct from other potato isolates.

In this study, evidence has been presented indicating that Mexican Fire disease is caused by PVY infection. The application of Koch’s Postulates has allowed for the establishment of a direct link between PVY infection and the development of necrosis in tomato leaves. Necrotic symptoms typically manifest around two weeks after the virus is detected. Initially, faint necrosis appears on the underside of leaves, often escaping notice without careful observation. Subsequently, the necrosis spreads to the upper side of the leaves. Unlike most virus diseases, initial symptoms tend to emerge in the middle leaves before progressing to the upper leaves. This results in the significant coverage of leaf blade area, giving rise to a distinctive burnt appearance. Importantly, while necrosis is predominantly observed in the middle to top leaves, the virus is also present in asymptomatic top leaves.

It is our belief that the challenge encountered in replicating necrotic symptoms through artificial inoculation in the greenhouse is likely attributed to the choice of cultivar for routine laboratory-based biological tests. Santa Clara is an open pollinated cultivar and is preferred due to the lower cost of the seeds; however, it does not exhibit leaf necrosis, despite its susceptibility to PVY infection (data not shown). In general, the observed symptoms were less pronounced than those documented in the field, a phenomenon previously documented by Costa et al. (1960), who extensively investigated PVY infections in diseased tomatoes. Notably, the cultivars Santy, Guará, and BS12 displayed high susceptibility to PVY infection and subsequent leaf necrosis.

The genome of Brazilian isolates of PVY found in tomato plants were grouped in a cluster with isolates classified as C strain, and all pepper isolates (Fig. 3). The two potato isolates present in this group were classified in the C strain, while the other O and N strains of potatoes included in the analysis were distantly grouped. It suggests that the Brazilian tomato PVY isolates are potentially part of the C strain, and that this group is prevalent in pepper plants. Based on the clear clustering of our isolates far from other isolates, it is expected that these viruses evolved in non-potato plants. Finally, we speculate that the Mexican Fire appeared as a consequence of the use susceptible cultivars that respond to PVY infection with necrosis starting in the median part of the plant.

In conclusion, our study underscores the critical importance of precisely identifying the etiological agent responsible for diseases such as Mexican Fire in tomato fields. Our findings highlight the significant contribution of old tomato and maize fields to the higher incidence rates of PVY, emphasizing the need for targeted management strategies in these areas. Notably, our observation of alatae aphids in yellow traps, despite the absence of apterous aphids in the monitored field, suggests their role as a relevant source of PVY, necessitating control measures outside the field. We observed a two-week interval from PVY infection to the onset of symptoms, with necrosis causing substantial damage to the plants. This necrosis may potentially be mistaken for symptoms caused by tospoviruses, underscoring the importance of accurate disease diagnosis. Urgent testing for resistance against PVY infection in commercial tomato cultivars is imperative to mitigate the impact of Mexican Fire disease on tomato production.

## Conceptualization

Alice Kazuko Inoue-Nagata; Methodology: all authors; Data and sample collection: Vivian dos Santos Lucena, José Luiz Pereira, Camila de Morais Rêgo Machado, Cristiano da Silva Rodrigues, Bernardo Ueno and Alice Kazuko Inoue-Nagata; Formal analysis: Vivian dos Santos Lucena, Erich Yukio Tempel Nakasu, Ivair José Morais and Alice Kazuko Inoue-Nagata; Writing - original draft preparation: all authors; Writing - review and editing: Vivian dos Santos Lucena, Erich Yukio Tempel Nakasu, Ivair José Morais, Camila de Morais Rêgo Machado, Cristiano da Silva Rodrigues and Alice Kazuko Inoue-Nagata; Funding acquisition and supervision: Alice Kazuko Inoue-Nagata

## Data Availability Statement

The nucleotide sequences obtained in this investigation have been deposited in the GenBank database under the following accession numbers: PVY-TOMNEC (OR497354), PVY-RNY1 (OR497355), and PVY-RNY2 (OR497356).

## Acknowledgements

The authors express gratitude to the grower João Generoso Caixeta Filho for granting access to the fields for our studies. Additionally, we extend our appreciation to Fabiano Ibraim Regis Carvalho from the Emater extension service for providing invaluable support during the field investigation. Special thanks are also due to Lucio Flávio Barbosa and Hamilton José Lourenço for their technical assistance in the greenhouse and laboratory settings.

## No conflict of interest exists

We wish to confirm that there are no known conflicts of interest associated with this publication and there has been no significant financial support for this work that could have influenced its outcome.

---

**Supplementary Table 1.** Three plants cv. Santy were evaluated for the presence of necrotic symptoms in each of the leaves and detection of PVY by ELISA.

| Leaf position | Plant 1 |  | Plant 2 |  | Plant 3 |  |
| --- | --- | --- | --- | --- | --- | --- |
|  | Symptom | ELISA | Symptom | ELISA | Symptom | ELISA |
| 1 | _* | - | - | + | - | + |
| 2 | - | + | - | + | - | + |
| 3 | - | + | + | + | - | + |
| 4 | - | + | + | + | - | + |
| 5 | - | + | + | + | + | + |
| 6 | + | + | + | + | + | + |
| 7 | + | + | + | + | + | + |
| 8 | + | + | - | + | + | + |
| 9 | + | + | + | + | + | + |
| 10 | - | - | + | + | + | + |
| 11 | - | - | - | - | + | + |
| 12 | + | + | - | - | + | + |
| 13 | + | + | - | - | - | + |
| 14 | - | + | - | - | - | - |
| 15 | - | + | - | - | - | - |
| 16 | - | + | - | - | - | - |
| 17 | - | - | - | - | - | - |
| 18 | X | X |  |  | - | - |
| 19 | X | X |  |  |  |  |
| 20 | X | X |  |  |  |  |
| 21 | X | X |  |  |  |  |

\*-: negative; +: positive; X: Eliminated.

